# Oral bioavailability and metabolism of hydroxytyrosol from food supplements

**DOI:** 10.1101/2022.11.16.516752

**Authors:** Cecilia Bender, Sarah Strassmann, Christian Golz

## Abstract

Table olives and olive oils are the main dietary sources of hydroxytyrosol (HT), a natural antioxidant compound that has emerged as a potential aid in protection against cardiovascular risk. Bioavailability studies with olive oils showed that HT is bioavailable from its free form and from conjugated forms like oleuropein and its aglycone. Still, its low dietary intake, poor bioavailability, and high inter-individual variability after absorption through the gastrointestinal tract hamper its full benefits.

In a randomized, controlled, blind, cross-over study, we investigated the impact of HT metabolism and bioavailability by comparing two olive-derived watery supplements containing different doses of HT (30.58 and 61.48 mg of HT/dosage, respectively); additionally, HT-fortified olive oil was used in the control group. To this aim, plasma and urine samples were evaluated in 12 healthy volunteers following the intake of a single dose of the supplements or fortified olive oil. Blood and urine samples were collected at baseline and at 0.5, 1, 1.5, 2, 4, and 12 hours after intake. HT and its metabolites were analyzed by UHPLC-DAD-MS/MS.

Pharmacokinetic results showed that dietary HT administered through the food supplements is bioavailable and biovailability increases with the administered dose. After intake, homovanillic acid, HT-3-*O*-sulfate, and 3,4-dihydroxyphenylacetic acid are the main metabolites found both in plasma and urine. The maximum concentrations in plasma peaked 30 minutes after intake.

Being the bioavailability of a compound a fundamental prerequisite for its effect, these results promise a good potential of both food supplements for the protection against oxidative stress and the consequent cardiovascular risk.

## 1. Introduction

Olive fruits (*Olea europaea L*.) contain several types of phenols mainly tyrosol (Tyr) derivates, phenolic acids, flavonols and flavones, which contents vary with the olive cultivar [1–2], the fruit maturity and time of harvest [3–4]. Hydroxytyrosol (HT; 3,4-dihydroxyphenyl-ethanol) is the main phenolic phytochemical occurring in olives [5] as a simple phenol or either esterified with elenolic acid to form oleuropein (Ole) aglycone. It shows potential antioxidant, anti-inflammatory, and health benefits mainly related with cardiovascular diseases [6–9]. Moreover, a health claim of olive oil phenolics is authorized in the EU, this indication states that a minimum daily intake of 5 mg of hydroxytyrosol and its derivatives protects low-density lipoprotein (LDL) cholesterol from oxidative damage [10–11]. In fact, it has been shown that olive oil phenolics in general, and HT and its derivatives in particular, bind to LDL cholesterol in humans [12] reducing its oxidation [13–16].

However, the olive phenolics relevant to health are not only contained in the oil but also in the water-soluble part of the olives. Indeed, HT is a polar molecule only slightly soluble in fats, a feature that hinders its passage through the lipid bilayer membrane in the small intestines.

Clinical evidence related to HT metabolism is based primarily on human studies conducted with olive oils containing varying concentrations of naturally present or added olive phenolic compounds [17–28]. Overall, these studies showed that despite the poor bioavailability of HT [17–18], it can be rapidly absorbed from the intestine [19], metabolised in the gut and liver, distributed and rapidly eliminated via the kidneys. Only a small fraction of free (unchanged) HT is detected in plasma or urine, as HT undergoes several changes through phase I / II metabolism. The main metabolites of the HT transformation result from the direct intestinal phase II conjugation (glucuronidation and sulfation), and from the enzymatic methylation-oxidation that gives rise to homovanillic acid (HVA). Furthermore, enzymatic oxidation of HT via alcohol-dehydrogenases and aldehyde-dehydrogenases gives rise to 3,4-dihydroxyphenylacetic acid (DOPAC), which can be further methylated via catecholmethyltransferase to form HVA [reviewed in 9, 29]. Additionally, and despite their lower concentration in olives, the absorbed Ole and Tyr can be metabolised into free-HT for absorption in the human body, thus increasing the concentration of HT in circulation [9, 29]. Pharmacokinetic data obtained with animal models also point out an extensive and fast uptake of HT, which is distributed in the body and detected dose-dependently in blood and urine as well as in different organs including kidneys, liver, heart and brain [25, 30–33].

Vegetation water is a by-product of olive oil processing, i.e., the aqueous part of the olive that is separated from the oil. Such liquid by-product is rich in polar phenols, and typically contains 98% of the total phenols from the olive fruit [34]. Olive phenolics are a promising functional ingredient for the prevention of cardiovascular disease and inflammatory events. However, in the field of food supplements, the absorption and bioavailability of the active ingredients in humans is an important issue that is often underestimated. And given that the bioavailability of HT and its derivates can be conditioned by the matrix with which they are administered [35–36], it is important to clarify whether its absorption, metabolism, and bioavailability occurs with a food matrix other than oil before claiming any beneficial implications for human health.

Our study investigates the bioavailability, disposition and dose response of HT in humans after intake of the proprietary food supplements, named *Oliphenolia* and *Oliphenolia bitter*. They are produced using the vegetation water generated during olive oil production, and contain mainly natural HT as simple phenol or esterified with elenolic acid to form Ole aglycone.

To this end, a randomized, controlled human trial was conducted with 12 healthy volunteers. Kinetics and bioavailability were determined analysing HT and Ole, as well as their metabolised oxidated-methylated, sulfated and glucuronidated forms, both in plasma and urine at 0.5 hr, 1 hr, 1.5 hr, 2 hr, 4 hr and 12 hr after taking 25 ml (1 flask) of the liquid supplement, and evaluated as a change from baseline. HT and its metabolites were analyzed by UHPLC-DAD-MS/MS using pure reference standards.

Results indicate that HT is dose-dependently absorbed after intake of the aqueous food supplements, it is metabolised mainly to HVA, HT-3-*O*-sulfate (HT-3-S), and DOPAC, and it is highly excreted in the urine.

These results promise a good potential of both food supplements for oxidative stress control *in vivo*. We hypothesized that the food supplements rich in natural HT may provide similar benefits on risk factors for cardiovascular disease to those provided by high phenolic extra virgin olive oils (EVOOs). Thus, further secondary outcomes of the present study [37] evaluated the potential antioxidant-health benefits of both food supplements, considering the postprandial kinetics of F2alpha-isoprostanes in urine and oxidized LDL in blood, as the HT from these supplements is expected to show positive effects on these markers of lipid peroxidation.

## 2. Materials and Methods

### 2.1 Standards and Reagents

HT, Ole, HVA, and DOPAC (all purity ≥ 98%) were from Cayman Chemical (Ann Arbor, MI, USA). Pure HT-3-S (98%) was supplied by ChemCruz (Huissen, the Netherlands); Tyr (purity ≥ 99,5%) was from Sigma-Aldrich (Taufkirchen, Germany). HT-3-G (purity ≥ 97%) was from Biozol (Eching, Germany); citric acid, phosphoric acid, and L(+)-ascorbic acid were from Roth (Karlsruhe, Germany). Water, acetonitrile, methanol, acetic acid and formic acid (all LC-MS-grade) were purchased from VWR Chemicals (Darmstadt, Germany).

Liquid food supplements *Oliphenolia bitter* and *Oliphenolia* and the EVOO were provided by Fattoria La Vialla S.A.S (Castiglion Fibocchi, Arezzo, Italy). HTEssence Hydroxytyrosol Liquid was kindly provided by Wacker Chemie AG (Munich, Germany).

### 2.2 Investigational Products (IPs)

The food supplements are derived from olive fruit (*Olea europaea L.*) vegetation water subjected to filtration and concentration. General composition of *Oliphenolia bitter* (hereafter referred to IP-1) and *Oliphenolia* (hereafter IP-2) is shown in table 1. IP-1 consists of a 94% concentrated vegetation water and 6% lemon juice (*Citrus limon L*. fructus). IP-2 is composed by 30% further concentrated vegetation water and 70% grape juice (*Vitis vinifera L*. fructus).

**Table 1.** Typical nutritional value of the food supplements, and content of hydroxytyrosol (HT) and derivatives (as free forms) in the food supplements used for the trial. Ole: oleuropein, Tyr: tyrosol, HVA: homovanillic acid, DOPAC: 3,4- dihydroxyphenylacetic acid. n.d.: not detectable; *:according to producer

| Nutritional value * (g/100 mL) | IP-1 | IP-2 |
| --- | --- | --- |
| Energy (kJ) | 75.00 | 555.00 |
| Fat | 0.00 | 0.05 |
| of which saturates | 0.00 | 0.00 |
| Carbohydrate | 4.17 | 32.03 |
| of which sugars | 1.05 | 27.52 |
| Protein | 0.20 | 0.34 |
| Salt | 0.028 | 0.019 |
| Potassium | 0.804 | 0.642 |
| **HT and derivatives (mg/L)** |  |  |
| HT | 1223 | 2459 |
| Ole | 1.65 | 2.99 |
| Tyr | n.d. | n.d. |
| HVA | n.d. | n.d. |
| DOPAC | n.d. | n.d. |
| Total polyphenols * | 10980 | 10100 |

Quantification of HT and derivatives in the food supplements was conducted as previously described [38]. The administrated dose (25 mL – 1 flask) of the food supplements were: 30.6 mg HT: 0.04 mg Ole for IP-1, and 61.5 mg HT: 0.07 mg Ole for IP-2. Additionally, EVOO was spiked with HT to reach a final concentration of 5.77 mg/20 g, concentration recognized to promote human health according to EFSA [11], and used in the control group. Free tyrosol, DOPAC and HVA were not detectable in any of the IPs. The sum of HT and Tyr in the fortified-EVOO was further measured after acid hydrolysis by ADSI GmbH (Innsbruck, Austria) via HPLC-UV and resulted equivalent to 12.19 mg (sum of HT + Tyr) per 20g of fortified EVOO.

### 2.3 Participants and study design

The present study has been registered at ClinicalTrials.gov (identifier: NCT04876261). The protocol was approved by the Ethics Commission of State Medical Association of Rheinland-Pfalz (Mainz, Germany). The clinical study was conducted as set out in the Declaration of Helsinki by *daacro GmbH & CO* at the Science Park Trier (Trier, Germany). Written informed consent was obtained from all participants.

Volunteers meeting the inclusion and exclusion criteria were recruited. Specific diet indications included to avoid consuming any olive derived products as well as alcohol and food supplements with hydroxytyrosol, vitamins, minerals, and antioxidants, at least 2-4 days before the first intake and during the whole study. Three days prior to and at each intervention, volunteers avoided moderate or intense physical activity. Volunteers underwent a wash-out period of 6 days between the interventions to avoid interference between the IPs.

Study design was randomized, single-blind, single-dose, three way cross-over, in which 12 healthy male volunteers ingested, after an overnight fast of at least 10 hrs, different concentrations of olive phenolics through the respective IP administrated with 200 mL of water.

One volunteer dropped out of the study after the completion of the first intervention period and was subsequently replaced, considering the data of this subject the sample size for the IP-2 group is 13.

### 2.4 Sampling

At each intervention visit, a baseline blood sample (9 mL) was collected immediately before administration of the IP. Further blood samples (9 mL each) were collected 0.5, 1, 1.5, 2,4 and 12 hr after the intake. EDTA-plasma samples were obtained, samples were stabilized with 10% V/V of 2 M aqueous citric acid.

At each intervention visit, a baseline urine sample was collected from –240 to 0 minutes before administration of the IP. Further urine samples were collected after the intervention from 0 to 30 min, 30 to 60 min, 60 to 90 min, 90 to 120 min, 120 to 240 min, and 4hr to 12hr. Urine samples were stabilized with 1.88 g/L of ascorbic acid and its volume was measured in order to be able to convert the proportion of excreted metabolites at each measurement point to the excreted volume in a comparable manner.

Stabilized plasma and urine samples were stored at −80 °C until analysis.

### 2.5 Sample processing

The method developed to purify the plasma and concentrate the HT metabolites was based on a previous research study [39]. Prior to analysis, plasma samples were thawed and mixed 1:1 with aqueous phosphoric acid (4%) to reduce phenolic-protein interactions. After centrifugation, the supernatant was plated on a 96 well Oasis HLB μElution SPE Plate from Waters (Eschborn, Germany) without conditioning. As washing solutions 200 μL of water and 200 μL of 0.2% acetic acid were added. The phenolic metabolites were eluted by 100 μl of acetonitrile + water (1+1) applied in two portions of 50 μL each.

Urine samples were thawed, filtrated through 0.20 μm regenerated cellulose filters (Macherey-Nagel, Düren, Germany) and immediately analyzed by UHPLC without further preparation.

Plasma standards were prepared in a control plasma which did not contain the tested compounds; urine standards were prepared in water. These reference standards (5.6 −0.02 mg/L) were prepared freshly for each measurement, and processed in the same way as the samples.

Peaks of HT metabolites were identified by comparison with reference compounds regarding retention time and MS^2^ spectra. For quantitation, a linear calibration equation was calculated from the peak area (MS^2^) vs concentration plot.

### 2.6 Analysis of HT and its metabolites

All samples were run in duplicate. Measurement was conducted by UHPLC-DAD-MS/MS, with an Acquity UPLC I-Class system coupled to a XEVO-TQS micro mass spectrometer (Waters, Milford, MA, USA). The instrument consisted of a sample manager cooled at 10 °C, a binary pump, a column oven, and a diode array detector (DAD) measuring at 280 nm for confirmation of the peaks. The column oven temperature was set at 40 °C. Eluent B was water with 0.1% formic acid, eluent A was acetonitrile with 0.1% formic acid, the flow was 0.4 mL/min on an Acquity BEH C18 RP column (50 mm × 2.1 mm, 1.7 μm particle size) combined with an Acquity BEH C18 precolumn (2.1 mm × 5 mm, 1.7 μm), both from Waters (Milford, MA, USA). The gradient started with 1% A for 3 min and raised linearly to 20% A within 1 min, then to 80% A within 1.3 min, then to 100 % A in 0.3 min and holding for 1.5 min as a washing step; back to 1% A within 0.2 min and equilibrating for 1 min. The injection volume was 5 μL.

The peaks were identified by MS/MS in negative mode. The source voltage was kept at 3.4kV, and the cone voltage was 77 V. The source temperature was set at 150 °C and the desolvation temperature at 400 °C with a desolvation gas flow of 800 L/h and a cone gas flow of 50 L/h. Data were acquired and processed using MassLynx 4.1 (Waters, Milford, MA, USA).

### 2.7 Data analysis

Each metabolite measurement was conducted in duplicate. The average values of concentration for each of the samples were calculated in Microsoft Excel version 16.0. For the raw statistics, classical statistical methods using median, mean, standard deviation and confidence intervals were used. Average values were further processed using GraphPad software (San Diego, California, USA) version 5.00 to represent in graphs and tables. Pharmacokinetic parameters maximum plasma concentration C_max_ (nmol/L), time to reach C_max_ (t_max_, h), and mean area under the concentration-time curve (AUC_0-12hr_)) were calculated from plasma concentrations, while cumulative concentrations were calculated for the urinary excretion. The data are expressed as mean ± standard error of the mean (SEM) unless otherwise indicated. Student’s unpaired t-test was performed to compare the results before and after ingestion for each test product.

In order to identify potentially hidden data correlations, artificial intelligence algorithms were applied by using MATLAB 2019 (Natick, Massachusetts, USA) software, including the machine learning and deep learning toolbox version 11.5. This includes the classification of the data using machine learning techniques (such as k-nearest neighbours, decision tree, and support vector machine), deep learning using deep neural networks as well as the analysis of the data using correlation matrix calculations.

## 3. Results

Table 2 shows the mass transitions used for the quantification and identification of HT and its metabolites as determined by fragmenting standard substances in the chromatograms and comparing these with the literature [40]. For HT-S and HT-G only the 3-*O*-standards were available, as these have the same mass transitions as their 4-*O*-isomeres, it was concluded that the second peak with the same mass transition detected in a similar retention time window is the 4-*O*-derivative.

**Table 2.**
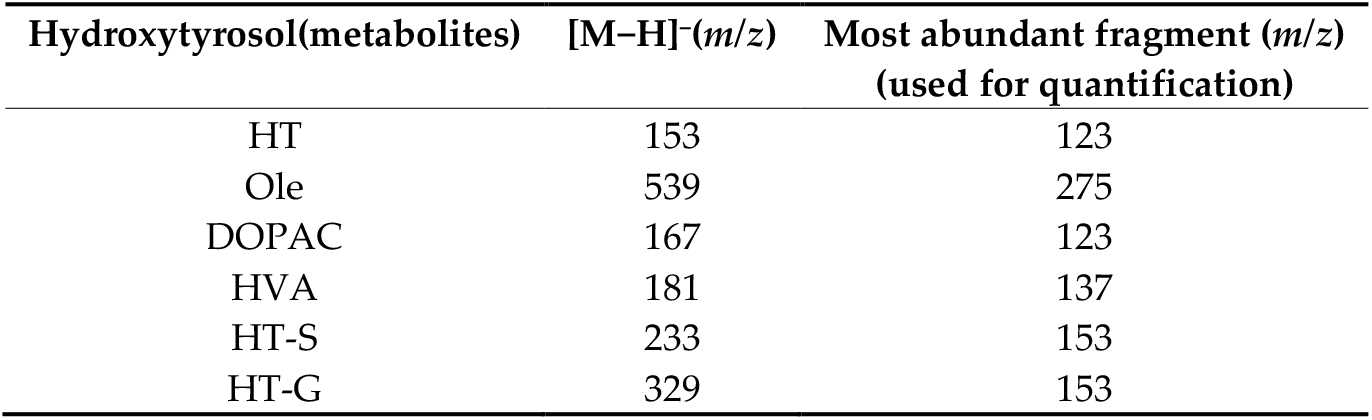
Mass transitions of hydroxytyrosol and its metabolites. HT: hydroxytyrosol, Ole: oleuropein, DOPAC: 3,4- dihydroxyphenylacetic acid, HVA: homovanillic acid, HT-S: hydroxytyrosolsulfates, HT-G: hydroxytyrosol glucuronides

| Hydroxytyrosol(metabolites) | [M-H]<sup>-</sup>(*m/z*) | Most abundant fragment (*m/z*)<br>(used for quantification) |
| --- | --- | --- |
| HT | 153 | 123 |
| Ole | 539 | 275 |
| DOPAC | 167 | 123 |
| HVA | 181 | 137 |
| HT-S | 233 | 153 |
| HT-G | 329 | 153 |

The HT metabolites were separated in a single UHPLC run. The respective limits of quantification (LOQ) were calculated using diluted reference solutions and the signal noise ratio of 10 (table 3).

**Table 3.** Limit of Quantification of HT and its metabolites in plasma and urine. Results are expressed as mean (in mg/L) ± SEM

|  | Plasma | Urine |
| --- | --- | --- |
| HT | 0.02 ±0.01 | 0.01 ±0.00 |
| Ole | 0.01 ±0.00 | 0.01 ±0.00 |
| HT-3-S | 0.05 ±0.01 | 0.05 ±0.00 |
| HVA | 0.08 ±0.02 | 0.14 ±0.08 |
| DOPAC | 0.25 ±0.04 | 0.09 ±0.03 |
| HT-3-G | 0.64 ±0.22 | 0.05 ±0.00 |

Detection of Tyr was interfered due to the co-elution of matrix components, consequently, the method was found to be unsuitable for quantification of Tyr in both, plasma and urine.

### 3.1 Plasma bioavailability

The non-metabolised forms of HT and Ole were almost undetectable in plasma after ingestion of the IPs. For the fortified EVOO group, the HT metabolites were almost undetectable in all subjects before and after the intake, with the exception of HVA that was detected (<LOQ) but quantifiable in two subjects before intake and 30 minutes after.

Table 4 illustrates the pharmacokinetic parameters from plasma concentrations of the main metabolites detected after the intake of IP-1 and IP-2. Although baseline plasma concentrations were below the LOQ, after ingestion of the food supplements the HT was detected primarily as HVA, followed by smaller amounts of HT-3-S and DOPAC. As expected, the mean areas under the concentration-time curves were higher for IP-2 than for IP-1, however, these differences are not significant (p>0.05) due to the large intra- and inter-individual variation observed, which is assumed to be mainly due to biological variability.

**Table 4.** Pharmacokinetic parameters from plasma concentration. Cmax: peak plasma concentrations expressed in nmol/L ± SEM; tmax: time (in minutes) to reach Cmax; AUC: mean area under the plasma concentration-time curve from 0 hr to 12 hr. n.s.: not significant difference

| IP | HT-3-S |  |  | HVA |  |  | DOPAC |  |  |
| --- | --- | --- | --- | --- | --- | --- | --- | --- | --- |
|  | C<sub>max</sub> | t<sub>max</sub> | AUC | C<sub>max</sub> | t<sub>max</sub> | AUC | C<sub>max</sub> | t<sub>max</sub> | AUC |
| IP-1 | 384.89<sup>n.s.</sup> ± 62.63 | 30 | 372<sup>n.s.</sup> | 868.87<sup>n.s.</sup> ± 111.86 | 30 | 1019<sup>n.s.</sup> | 301.07<sup>n.s.</sup> ± 134.43 | 30 | 169<sup>n.s.</sup> |
| IP-2 | 406.28<sup>n.s.</sup> ± 62.68 | 30 | 443<sup>n.s.</sup> | 948.76<sup>n.s.</sup> ± 160.19 | 30 | 1672<sup>n.s.</sup> | 466.96<sup>n.s.</sup> ± 147.68 | 30 | 247<sup>n.s.</sup> |

Comparing the kinetic of plasma concentration of the major metabolites HVA peaks at 30 minutes and progressively decreases over the next 3.5 hours (Figure 1); HT-3-S reached the maximum concentration 30 min after intake and strongly decreased within 2 hr after administration. Of note, HVA and HT-3-S, were measurable from 0.5 to 2 hr at the lower HT-dose and from 0.5 to 4 hr at the higher dose. DOPAC showed the maximum concentration at 30 minutes after the intake of HT through the food supplements followed by a marked decrease until reaching values close to the LOQ one hour after ingestion.

**Figure 1.**
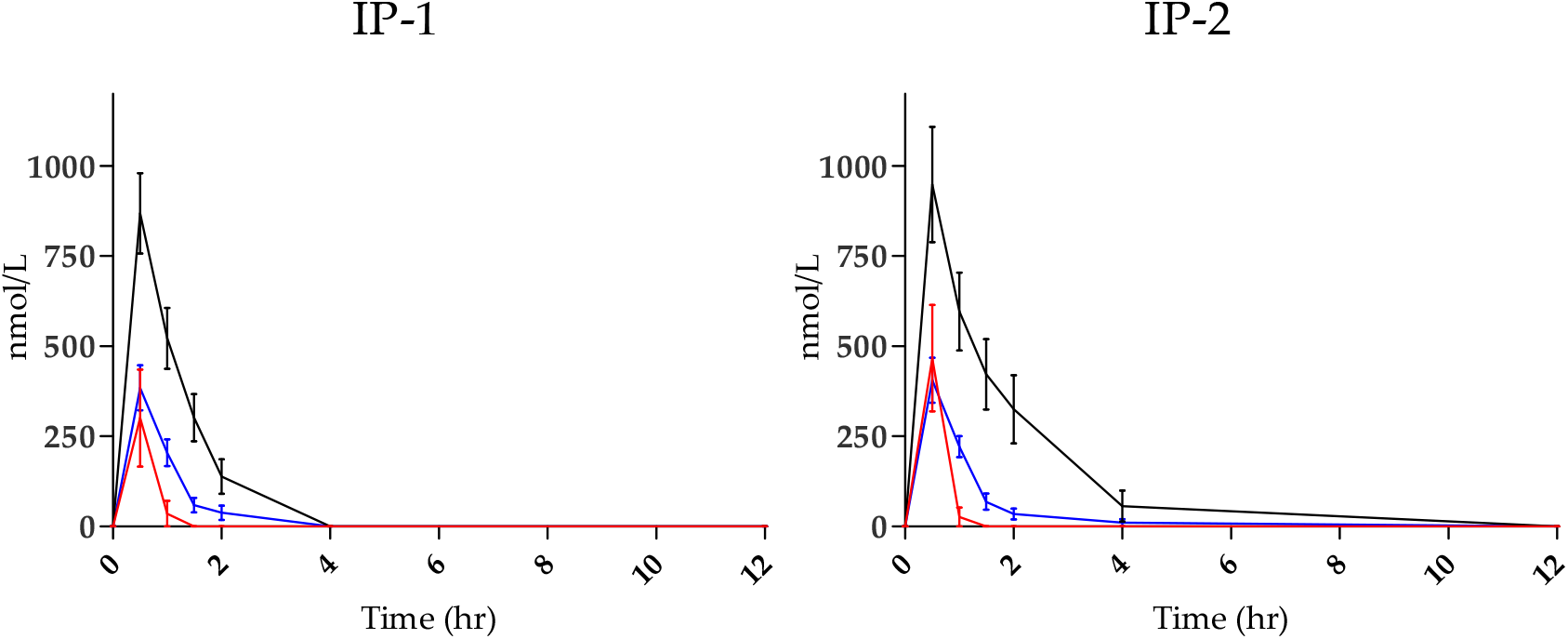
Mean plasma concentration profile of main HT metabolites before (0 hr) and after intake of IP-1 (n=12) and IP-2 (n=13). HVA: black lines; HT-3-S: blue lines; DOPAC: red lines. Bars: SEM.

Among the other HT metabolites studied, HT-4-S was undetectable in plasma from all groups, while the glucuronide conjugates were poorly present: HT-3-G was mainly detected in trace amounts (<LOQ) after all the interventions, but measured in one subject at 0.5 and 1 hr after intake of the highest HT-dose (IP-2), peaking at 1 hour after intake (Cmax: 8.73 nmol/L). HT-4-G was detected (<LOQ) only in few samples after intake of the food supplements but not after intake of the fortified EVOO.

### 3.2 Urinary excretion

The unmetabolised form of Ole was almost undetectable in the urine before and after ingestion of the respective IPs. Free (unchanged) HT was excreted within an hour and in small amounts after the ingestion of IP-1 and IP-2 (0.0004 μmole ± 0.0010 and 0.0024 μmole ± 0.0009, respectively; not significant difference between groups), but not after EVOO intake (p < 0.05 vs IP-2).

A dose-dependent concentration of all the HT-metabolites analysed was recorded following the IPs administration (Table 5). Moreover, the three main metabolites quantified in plasma were extensively excreted in urine after intake in all the interventional groups (Figure 2). For instance, after the ingestion of the HT through the aqueous food supplements, the main metabolite excreted was DOPAC, followed by HVA (p < 0.05), and HT-3-S (p < 0.001), while after EVOO intake the HVA ranked first but was excreted in a similar extent to DOPAC and HT-3-S (not significantly different at p < 0.05 level).

**Table 5.** Mean cumulative (0-12 hr) excretion of HT-metabolites (micromoles) ± SEM in urine. Significance vs EVOO was as follows: n.s: not significant different, *: significant at p < 0.001 level.

| IP | n | HVA | DOPAC | HT-3-S | HT-4-S | HT-3-G | HT-4-G |
| --- | --- | --- | --- | --- | --- | --- | --- |
| IP-1 | 12 | 36.01* ± 20.08 | 44.31* ± 17.52 | 16.58* ± 6.0 | 5.32<sup>n.s</sup> ± 16.23 | 0.46* ± 0.35 | 0.15<sup>n.s</sup> ± 0.10 |
| IP-2 | 13 | 46.16* ± 5.37 | 59.74* ± 3.00 | 20.05* ± 1.55 | 0.16<sup>n.s</sup> ± 0.05 | 0.87* ± 0.13 | 0.41* ± 0.10 |
| EVOO | 12 | 4.84 ± 3.93 | 4.53 ± 1.76 | 2.57 ± 1.17 | 0.05 ± 0.19 | 0.06 ± 0.09 | 0.01 ± 0.02 |

**Figure 2.**
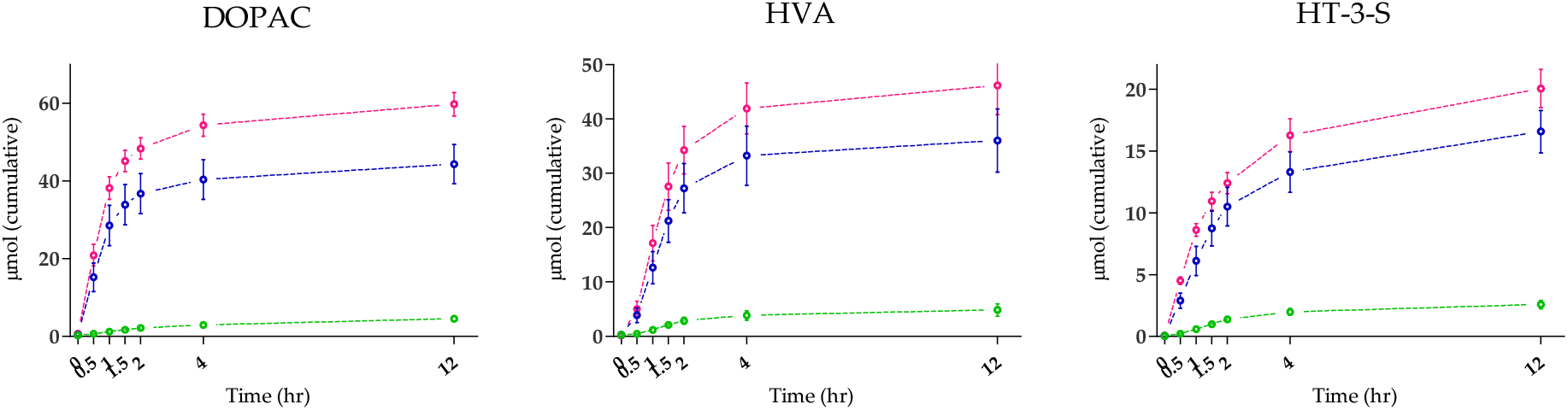
Mean urinary excretion of main HT metabolites quantified before (0 hr) and after intake of the investigational products. Blue lines: IP-1 (n=12), pink lines: IP-2 (n=13), green lines: EVOO (n=12). Bars: SEM.

The HT-4-S and the glucuronide conjugates were measured after intake in all the interventional groups, but in smaller concentrations than HVA, DOPAC and HT-3-S. Unlike IP-1, the EVOO and IP- 2 groups, containing the lowest and highest dose of HT, respectively, showed a predominant detection of HT-glucuronides over HT-4-S (not significant for EVOO, and p < 0.01 for IP-2).

The mean HT excretion calculated from the accumulated amounts was estimated at 59.6% and 35.8% of the total intake for IP-1 and IP-2, respectively, and 27.6% when administered with HT-enriched EVOO.

## 4. Discussion

The two food supplements studied here are based on ingredients of vegetable origin, mainly olive concentrate resulting from the aqueous by-product generated during olive oil production, and thus contain HT as the main bioactive. In addition, they contain lemon juice (IP-1) or grape juice (IP-2) concentrates. Despite olive phenolics being a promising functional ingredient, the bioavailability of HT and its derivates could be conditioned by the matrix with which they are administered [35–36]. Therefore, before claiming beneficial properties for human health is of key importance to verify the absorption, metabolism, and bioavailability of the active ingredient in the context of the food matrix that contains it.

In line with other studies conducted with olive oil and pure HT [12, 24–25, 41–43], our results show that HT administered together with the aqueous food supplements is absorbed from the intestinal tract in varying amounts and rapidly metabolised to phase I and phase II metabolites.

HT metabolites, which were undetected in plasma in the fasting state, were rapidly cleared from plasma in the postprandial phase (Cmax 30 min, complete clearance 2–4 hours) and excreted in the urine. The free (unchanged) forms of HT and Ole were almost undetectable in most plasma and urine samples, both before and after the intake of the products by the oral route and regardless the IPs, this result is in line with data in the literature [12, 20].

Similarly to previous studies reporting a dose-dependent absorption of HT contained in administered oil [17, 25, 41–43], the mean of HT metabolites detected in plasma (as the sum of all quantifiable metabolites) correlated with the ingested dose of HT, being higher for IP-2 than IP-1, although the difference did not reach statistical significance, and significantly higher than the fortified-EVOO (p < 0.05). A large inter-individual variability in the absorption of HT was observed, which has also been reported previously [12, 16, 19, 22, 36, 44].

The absolute amount of HT in urine (as the sum of all quantifiable metabolites in 12 hr) correlated with the dose administered, that is, fortified-EVOO < IP-1 < IP-2. However, the excreted percentage of the total ingested HT was as follows: fortified-EVOO < IP-2 < IP-1, overall the excretion ranged from 28 to 60 %, which is consistent with previous published data [25]. Contrary to what is reported in the literature [12, 35–36], the excretion of the HT administered as a natural component of an aqueous matrix can exceed that of the oily matrix. These results underline the importance of dietary factors influencing bioavailability, such as the possible presence of effectors (positive or negative) in the food matrix, the difference in phenolics, water or fat contents, all factors that can influence the absorption of HT itself. Similarly, synergistic interactions with other phenolic components of food matrices cannot be excluded [45].

The highest average concentrations of HT-metabolites in plasma (as the sum of all the quantifiable metabolites) were found 30 min after intake of the food supplements, being significantly different to the intake of EVOO, which only rarely showed levels above the quantification limit. HT is highly metabolised mainly to HVA, HT-3-S, and DOPAC; these metabolites (free forms) could often be detected in plasma samples from the food supplement groups and especially immediately after ingestion, but only very rarely in the fortified-EVOO group. Moreover, after ingestion of both supplements DOPAC and HVA reach the maximum plasma concentration at 30 minutes, but DOPAC concentration was lower and its elimination faster than that of HVA, thus is likely due to DOPAC’s enzymatic transformation into HVA via catecholmethyl transferase [17, 30].

Our findings indicate that regardless of the food matrix (i.e. watery or oily), the free forms of HVA and DOPAC, which result from the oxidation and/or methylation of dietary HT, contribute to a greater extent to the metabolisation of HT in the human body. To date, few human studies have looked for the free forms of these metabolites after HT ingestion. Earlier studies conducted with olive oil observed an increase of total HVA (after hydrolysis) in 24 hr pooled urine [17] and despite the high basal urinary concentration of HVA registered in all volunteers. Another human study conducted after the intake of ~175 mg of pure HT (99.5 %) in aqueous solution reported the excretion of DOPAC-glucuronide and HVA in 24 hr pooled urine, but not in plasma samples [12]. Overall our results are in line with these previous findings, and in particular with Kontouri and colaborators, whom addressed the conversion of HT into DOPAC, HVA, and homovanillic alcohol (both free and total) in humans after the ingestion of olives (fruit), these researchers reported that HT is excreted in urine and plasma mainly as HVA and DOPAC [24].

Among the phase II conjugates, sulfated metabolites (particularly HT-3-S) is preferred over glucuronidated ones. These results are in line whith previous studies showing that sulfation at position −3 is preferred to −4 [31, 46], and that sulfates are mainly detected after intake of high HT doses while glucuronates at low dosages [27, 31].

Overall, our results show that after ingestion of both aqueous dietary supplements, HT is absorbed and highly metabolised into phase I and phase II metabolites. These results promise a good potential of both food supplements for the protection against oxidative stress in the prostprandial phase; further secondary outcomes of the clinical trial are the potential antioxidant-health benefits of both food supplements, considering the postprandial kinetics of F2alpha-isoprostanes in urine and oxidized LDL in blood, as the HT from these supplements is expected to show positive effects on these markers of lipid peroxidation.

## Author Contributions

CB: Conceptualization, investigation, data analysis, original draft writing; SS: formal analysis, methodology; C.G.: data review with AI; CB, SS: review and editing. All authors have read and agreed to the published version of the manuscript.

## Institutional Review Board Statement

The study was conducted in accordance with the Declaration of Helsinki, approved by the Ethics Committee of State Medical Association of Rheinland-Pfalz (Mainz, Germany), and registered at ClinicalTrials.gov (identifier: NCT04876261).

## Informed Consent Statement

Informed consent was obtained from all subjects involved in the study.

## Acknowledgments

The authors would like to acknowledge the University Hospital Bonn (Germany) for kindly providing plasma samples for the preparation of standard curves in plasma, and Mrs. Eva Rogel (Institut Kurz GmbH) for her technical support.

